# Epilepsy-Associated SCN2A-L1342P Mutation Drives Network Hyperexcitability and Widespread Transcriptomic Changes in Human Cortical Organoids

**DOI:** 10.1101/2025.08.18.670956

**Authors:** Maria I. Olivero-Acosta, Morgan Robinson, Zhefu Que, Zaiyang Zhang, Hope Elizabeth Harlow, Vinayak Shankar, Seoyong Hong, Muhan Wang, Conrad M. Otterbacher, Hina Kadono, Manasi Halurkar, Harish Kothandaraman, Nadia Lanman, Trang Nguyen, Kyle Wettschurack, Benjamin Zirkle, Layan Yunis, Ningren Cui, Xiaoling Chen, Jingliang Zhang, Jiaxiang Wu, William C. Skarnes, Chongli Yuan, Feng Guo, Yang Yang

## Abstract

**Objective:** *SCN2A* pathogenic mutations, such as the recurrent heterozygous Nav1.2-L1342P, are monogenic causes of epilepsy. In this human-induced pluripotent stem cell model system, we aim to investigate the molecular and cellular mechanisms underlying the SCN2A-L1342P-associated pathology.

**Methods:** Using a human male iPSC reference line (KOLF) carrying the Nav1.2-L1342P mutation, we generated 3D cortical organoids for functional studies. Patch clamp and multi-electrode array (MEA) recordings, immunocytochemistry, and RNA sequencing were used to characterize the disease phenotypes.

**Results:** Nav1.2-L1342P organoid neurons displayed increased intrinsic excitability, and amplified excitatory post-synaptic currents, which are consistent with an increase in excitatory synapse formation revealed by PSD95/SYN1 immunostaining. Moreover, elevated network firing activity, as demonstrated by MEA, indicates a pronounced network hyperexcitability. Transcriptomic profiling of organoids carrying the Nav1.2-L1342P mutation further revealed significant alterations in synaptic, glutamatergic, and developmental pathways.

**Significance:** Our findings demonstrate that the Nav1.2-L1342P mutation drives a multifaceted disease phenotype, including network hyperexcitability and disruption of pathways related to neuronal and synaptic functions. These results advance our understanding of SCN2A-related Developmental and Epileptic Encephalopathy (DEE), laying a foundation for personalized interventions.

**Key Points:**

- **Key Point 1**: Human neurons carrying epilepsy-causing *SCN2A*-L1342P display intrinsic hyperexcitability in cortical organoids.
- **Key Point 2**: Synaptic transmission is enhanced in organoids carrying the *SCN2A*-L1324P mutation, consistent with the presence of elevated excitatory synapses, which leads to increased network hyperexcitability.
- **Key Point 3:** Transcriptomics analysis reveals that the *SCN2A*-L1342P mutation causes significant differential changes in forebrain developmental genes, synaptic and glutamatergic signaling pathways, and enhances cellular senescence and apoptosis.

## Introduction

The *SCN2A* gene, located on chromosome 2 (2q24.3), encodes the alpha subunit of the neuronal voltage-gated type II sodium channel Na_V_1.2, which plays a key role in action potential initiation and propagation (Pérez-Palma et al., 2020; Reynolds et al., 2020). Neuronal Na_V_1.2 is mainly expressed in the axonal initial segment (AIS) and somato-dendritic compartments of glutamatergic excitatory and other principal neurons (Kruth et al., 2020; Sanders et al., 2018; Spratt et al., 2021).

Mutations in SCN2A, particularly gain-of-function (GoF) variants, are a known cause of early infantile developmental epileptic encephalopathy (EIEE) (Sanders et al., 2018). Among these, the recurrent Na_V_1.2-L1342P (p.Leu1342Pro) mutation, resulting from a Leucine (L) to Proline (P) substitution within domain III of the sodium channel, has been identified in at least five patients worldwide (Crawford et al., 2021; Hackenberg et al., 2014; Matalon et al., 2014; Wolff et al., 2017). Biophysical studies have shown that the L1342P variant increases both window current and current density, consistent with a GoF phenotype (Que et al., 2025; Que et al., 2021). Clinically, affected individuals present with severe intellectual disability, hypsarrhythmia, infantile spasms, epileptic spasms, and poor eye contact, beginning within 6 months of birth (Crawford et al., 2021). Additional features include transient choreatic movement disorders and hypersomnia (Hackenberg et al., 2014). Brain atrophy has also been reported in four out of five individuals with the Na_V_1.2-L1324P variant, suggesting a potential mutation-specific effect on brain structural development (Crawford et al., 2021; Miao et al., 2020). Conventional treatments such as levetiracetam, vigabatrin, and prednisolone offer limited clinical benefit (Hackenberg et al., 2014; Matalon et al., 2014). Despite these findings, the molecular and cellular mechanisms by which the Na_V_1.2-L1342P mutation disrupts neuronal function remain poorly understood.

To study the molecular and cellular mechanisms underlying these complex phenotypes, we employed cortical organoids: human induced pluripotent stem cell (hiPSC) derived 3D self-organizing aggregates that resemble the dorsal forebrain (Revah et al., 2022). Cortical organoid models exhibit a neuroectoderm-like epithelium that matures over time, giving rise to cortical neurons (Yoon et al., 2019). At later stages of development, they show a heterogeneous cellular composition, predominantly comprising glutamatergic neurons with characteristics of both deep and superficial layers (e.g., CTIP2 and TBR1-positive neurons). These neurons form layer-like structures, establish synaptic connections, and display robust electrical activity, making them physiologically relevant models for electrophysiological studies (Sloan et al., 2017; Yoon et al., 2019).

In the present study, we established a 3D cortical organoid model to further investigate the molecular, cellular, and network consequences of the epilepsy-associated Nav1.2-L1342P mutation. Electrophysiological analyses of our 3D model using patch-clamp recordings revealed that Na_V_1.2-L1342P neurons exhibit enhanced repetitive action potential firing, increased intrinsic neuronal excitability, and elevated network-level neuronal activity. This hyperactive profile is further supported by a pronounced increase in excitatory post-synaptic currents (EPSCs) and a heightened number of excitatory synapses. In parallel, RNA sequencing of cortical organoids with the Na_V_1.2-L1342P mutation revealed significant dysregulation in synaptic, glutamatergic, neurodevelopment-related, and apoptotic/senescent transcriptional pathways, suggesting broad alterations in gene expression. These findings suggest that the Na_V_1.2-L1342P mutation drives not only electrophysiological changes but also widespread transcriptional disruptions. Taken together, our findings indicate that the molecular etiology of this GoF SCN2A-L1342P genetic mutation involves a convergence of hyperexcitability and multiple disrupted signaling pathways. These findings provide insights into the molecular and cellular mechanisms underlying SCN2A gain-of-function seizures and the abnormal brain structure observed in patients with this mutation.

## Results

### The Nav1.2-L1342P Mutation Enhances Intrinsic Excitability of HiPSC-Derived Cortical Organoid Neurons

We previously demonstrated that 2D cortical neuron monolayers carrying the Nav1.2-L1342P mutation exhibit a GoF phenotype (Que et al., 2021) and increased glutamate release (Que et al., 2025) compared to controls. However, 2D models lack the three-dimensional tissue architecture and network complexity required to fully capture the dynamic electrophysiological interactions of the brain. In this study, we utilized a 3D brain organoid model to investigate how the Nav1.2 L1342P mutation influences neuronal excitability at both single-cell and network levels, enabling us to explore the impact on synaptic connectivity and pathological firing patterns in a more physiologically relevant system. Three-dimensional cortical organoids **(Figure 1)** were generated from hiPSCs using a dual-SMAD inhibition medium protocol (Miura et al., 2022), supplemented with small molecules to induce cortical neuron fate **(Figure 1A-B)**. On day 30, we identified high proportions of neural progenitor cell populations positive for SOX2, which also had proliferative capacity, as seen by immunocytochemistry of the Ki67 proliferation marker **(Figure 1C)**. At day 120 of maturation, differentiated cortical organoids contain NEUN+ neurons **(Figure 1D)** and TBR1+ cortical neurons **(Figure 1E)**, which arranged themselves in a layer-like fashion. MAP2 staining was also found throughout the organoids **(Figure 1E),** demonstrating widespread neuronal architecture. These data provide evidence that the hiPSCs carrying Nav1.2-L1342P and their corresponding isogenic control can be differentiated into cortical organoids.

**Figure 1.**
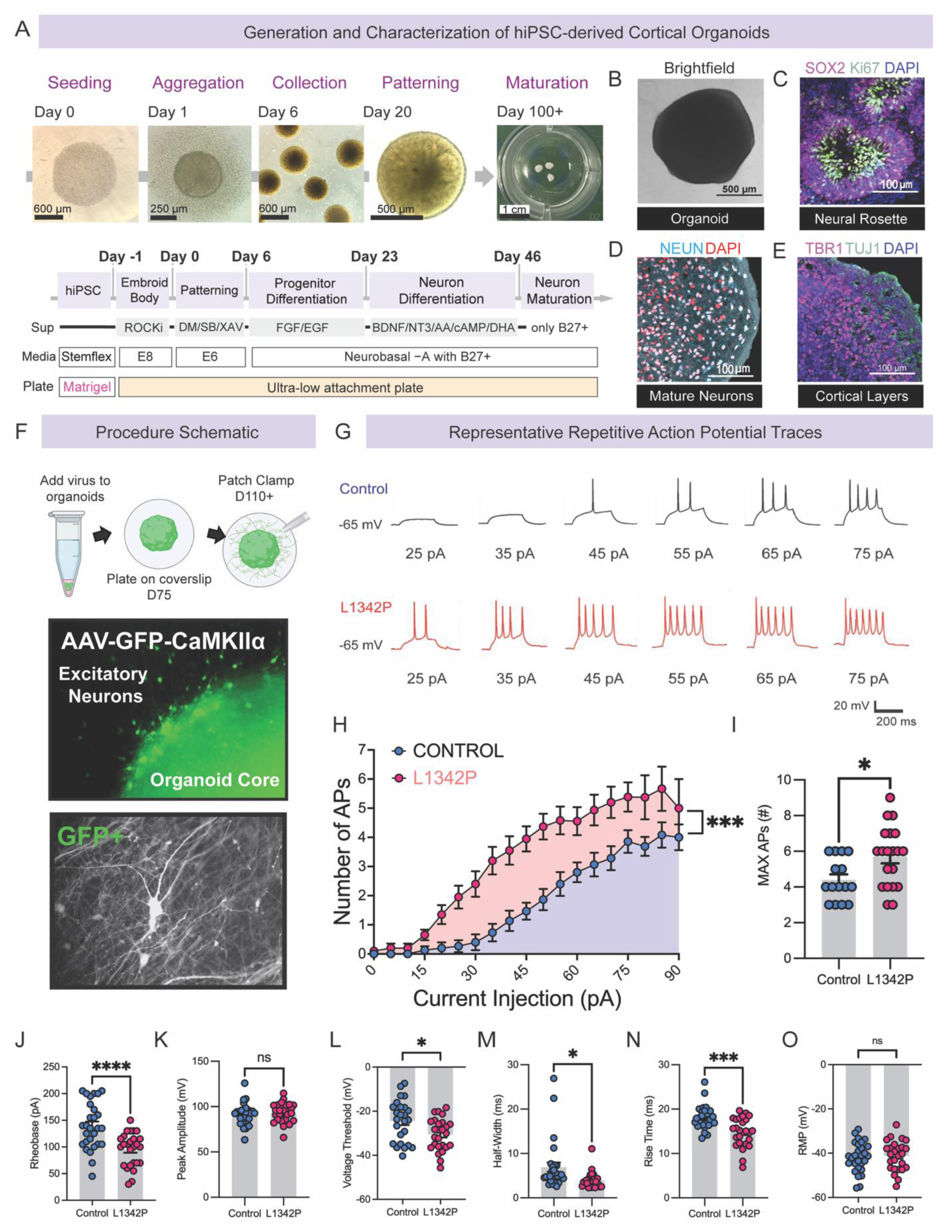
The Nav1.2-L1342P mutation enhances the repetitive firing of neurons in hiPSCs-derived cortical organoid. **(A)** Experimental schematic for the generation of hiPSC-derived cortical organoids. **(B)** Brightfield image of a cortical organoid in suspension. **(C)** Day 30 cortical organoid immunocytochemistry for neural progenitor marker SOX2, proliferation marker Ki67, and cell nuclei stain DAPI (blue). Neural rosettes are self-organizing, multi-cellular assemblies that hold neural progenitor cells (NPCs), radially organized in a central lumen resembling an embryonic neural tube, and appear as distinct, flower-like patterns. **(D)** 120-day-old cortical organoids express mature neuron marker NEUN (cyan). **(E)** 120-day-old cortical organoids express the cortical marker TBR1 (magenta), arranged in a layer-like formation, and MAP2 (green). **(F)** Preparation of hiPSC-derived cortical organoids for patch clamp experiments. 75-day-old organoids are exposed to a CamKIIa-GFP virus that labels excitatory neurons green and is plated onto coverslips. Over time, neurites emanate from the cortical organoid and become available to patch after day 111 of organoid development. **(G)** Representative sustained AP firings from hiPSC-derived Nav1.2-L1342P (red) organoid cortical neurons or controls (blue). **(H)** The plot shows the AP number per epoch in response to graded inputs from 0 to 90 pA current injections (400-ms duration). **(I)** The maximum number of action potential firings at increasing current injections was amplified in the Nav1.2-L1342P cortical organoid neurons. **(J)** The rheobase was decreased in Nav1.2-L1342P cortical organoid neurons. **(K)** The peak amplitude was unchanged between conditions. **(L)** The voltage threshold was reduced in Nav1.2-L1342P cortical organoids. **(M)** The half-width was reduced in Nav1.2-L1342P cortical organoids. **(N)** The rise time was decreased in Nav1.2-L1342P cortical organoids. **(O)** The resting membrane potential (RMP) was unchanged between conditions. Data are reported as mean ± error (S.E.M). Each dot corresponds to one neuron. Data in C was analyzed by *Repeated-measures two-way ANOVA*, ****p < 0.001, and data in D with an Unpaired Student’s t-test; *p < 0.05. Data in E-L was analyzed using the *Unpaired Student’s t-test*. * p < 0.05; ** p < 0.01; ****p < 0.001.

To investigate how the Nav1.2-L1342P mutation impacts both neuronal repetitive firing and the intrinsic properties of individual action potentials (APs), we performed whole-cell current-clamp. Cortical organoids were cultured 75–80 days, transduced with AAV-CaMKIIa-EGFP virus (for the identification of excitatory cortical neurons), plated onto PLO-Laminin-coated coverslips, and further cultured to 110–150 days before patch clamp recordings **(Figure 1F)**. Pyramidal-shaped neurons expressing EGFP were selected for patch-clamp analysis. We reveal a significant difference in repetitive firing between neurons carrying the Nav1.2 L1342P mutation and isogenic controls, using a prolonged 400-ms current stimulus ranging from 0 to 90 pA in 5-pA increments. From the 25 to 75 pA stimuli, the Nav1.2-L1342P neuron fired two to seven APs in response to increasing current steps **(Figure 1G, top)**. In contrast, zero or at most four APs were elicited in isogenic control neurons **(Figure 1G, bottom)**. Overall, the Nav1.2-L1342P neurons fired significantly more APs than controls in the 0–90 pA increasing current injections (*Repeated-measures two-way ANOVA analysis*, **Figure 1H**). Notably, the maximum number of AP firings that could be triggered by current injection was also significantly higher in Nav1.2-L1342P neurons, showing a 29.5% increase over control neurons (Control: 4.400 ± 0.3055 APs, n = 15 neurons, two clones across three differentiations; L1342P: 5.700 ± 0.3777 APs, n = 20 neurons, three clones across five differentiations, *\*p* = 0.0158, *Unpaired Student’s t-test,* **Figure 1I**). Our data suggests that the L1342P mutation significantly increased repetitive firing, indicating a hyperexcitable phenotype in the cortical organoid model.

While repetitive firing evaluates the neuronal response to sustained stimuli, examining the intrinsic properties of individual action potentials (APs) provides insights into how the mutation alters the biophysical properties of each AP, such as threshold, amplitude, and kinetics. Cortical neurons were held at –65 mV and depolarized with current injections in 5-pA increment steps, using a 20ms impulse, across 0-250pA current injections. Quantitatively, we found a 31.03% reduction in the rheobase of the Nav1.2-L1342P cortical organoid neurons compared to controls, indicating that less current is needed to elicit an action potential (AP) in the mutant condition, making them hyperexcitable (Control: 139.3 ± 8.244 pA, *n* = 28 neurons, two clones across five differentiations; L1342P: 95.19 ± 6.161 pA, *n* = 26 neurons, three clones, across seven differentiations; \*\*\*\**p*<0.0001, *Unpaired Student’s t-test*, **Figure 1J**).

Interestingly, we found that the AP amplitude did not differ between control and Nav1.2-L1342P neurons (Control: 90.48 ± 2.315 mV, *n* = 28 neurons, two clones across five differentiations; L1342P: 92.85 ± 2.087 mV, n = 26 neurons, three clones, across seven differentiations; ns *p* = 0.4346, *Unpaired Student’s t-test,* **Figure 1K**). We further analyzed the voltage threshold of spiking, as sodium channel dysfunction is likely to influence this parameter. Our results revealed that Nav1.2-L1342P neurons had a significantly lower voltage threshold, indicating a higher probability of AP firing at a more negative membrane potential (Control: –24.25 ± 1.888 mV, *n* = 25 neurons, two clones across five differentiations; L1342P: –30.53 ± 1.419 mV, *n* = 26 neurons, three clones, across seven differentiations; \**p* = 0.0127, *Unpaired Student’s t-test,* **Figure 1L**).

The Nav1.2-L1342P neurons also increased the kinetics of action potential (AP) firing. We found a statistically significant decrease in AP (spike) half-width in Nav1.2-L1342P cortical organoid neurons compared to controls, indicating a shorter AP duration (Control: 6.904 ± 1.055, n = 28 neurons, two clones across five differentiations; and L1342P: 3.942 ± 0.3562, n = 26 neurons, three clones, across seven differentiations; *p = 0.0128, *Unpaired Student’s t-test*, **Figure 1M**). Additionally, the AP rise time was reduced in the Nav1.2-L1342P neurons compared to controls, indicating a faster action potential initiation (Control: 18.12 ± 0.4984 ms, *n* = 28, two clones across five differentiations; L1342P: 14.80 ± 0.6431 ms, *n* = 26 neurons, three clones, across seven differentiations; \*\*\*\**p* = 0.0001, *Unpaired Student’s t-test,* **Figure 1N**). We also measured the resting membrane potential (RMP) to determine whether the Nav1.2-L1342P mutation affects this property. However, the RMP remained unchanged (**Figure 1O**).

### Nav1.2-L1342P Cortical Organoid Neurons Show Enhanced Excitatory Synaptic Transmission and Elevated Synapse Formation Compared to Controls

In addition to increased intrinsic excitability, spontaneous excitatory post-synaptic currents (sEPSCs) mediate synaptic transmissions that contribute to network excitability (Wu et al., 2022). To this end, we evaluated sEPSC in both control cortical organoids **(Figure 2A, top)** and those carrying the Nav1.2-L1342P variant **(Figure 2A, bottom)**. We found that the sEPSC frequency was markedly increased in the Nav1.2-L1342P organoids compared to controls (Control: 1.737 ± 0.3648, *n* = 17 neurons, three clones across five differentiations; L1342P: 5.142 ± 0.9129, *n* = 13 neurons, three clones across three differentiations; \*\**p* = 0.0021, *Mann-Whitney U test,* **Figure 2B**). The sEPSC amplitude of Nav1.2-L1342P cortical organoid neurons was significantly higher than that of controls (Control: 14.15 ± 0.6928, *n* = 17 neurons, three clones across five differentiations; L1342P: 17.79 ± 1.059, *n* = 13 neurons, three clones across three differentiations; \*\**p* = 0.0100, *Mann-Whitney U test*, **Figure 2C**), indicating a greater magnitude of synaptic events in the mutant neurons. Additionally, the EPSC inter-event-interval was reduced in the Nav1.2-L1342P cortical organoids compared to controls (Control: 1059 ± 193.1, *n* = 17 neurons, three clones across five differentiations; L1342P: 345.1 ± 94.28, *n* = 13 neurons, three clones across three differentiations; \*\**p* = 0.0021, *Mann-Whitney U* test, **Figure 2D**), indicating that the time between each synaptic event is shorter; therefore EPSC events occur more frequently. These findings suggest that the Nav1.2-L1342P cortical organoids have enhanced synaptic transmission compared to controls. These results support our previously reported finding thatglutamate release was increased due to the L1342P mutation (Que et al., 2025).

**Figure 2.**
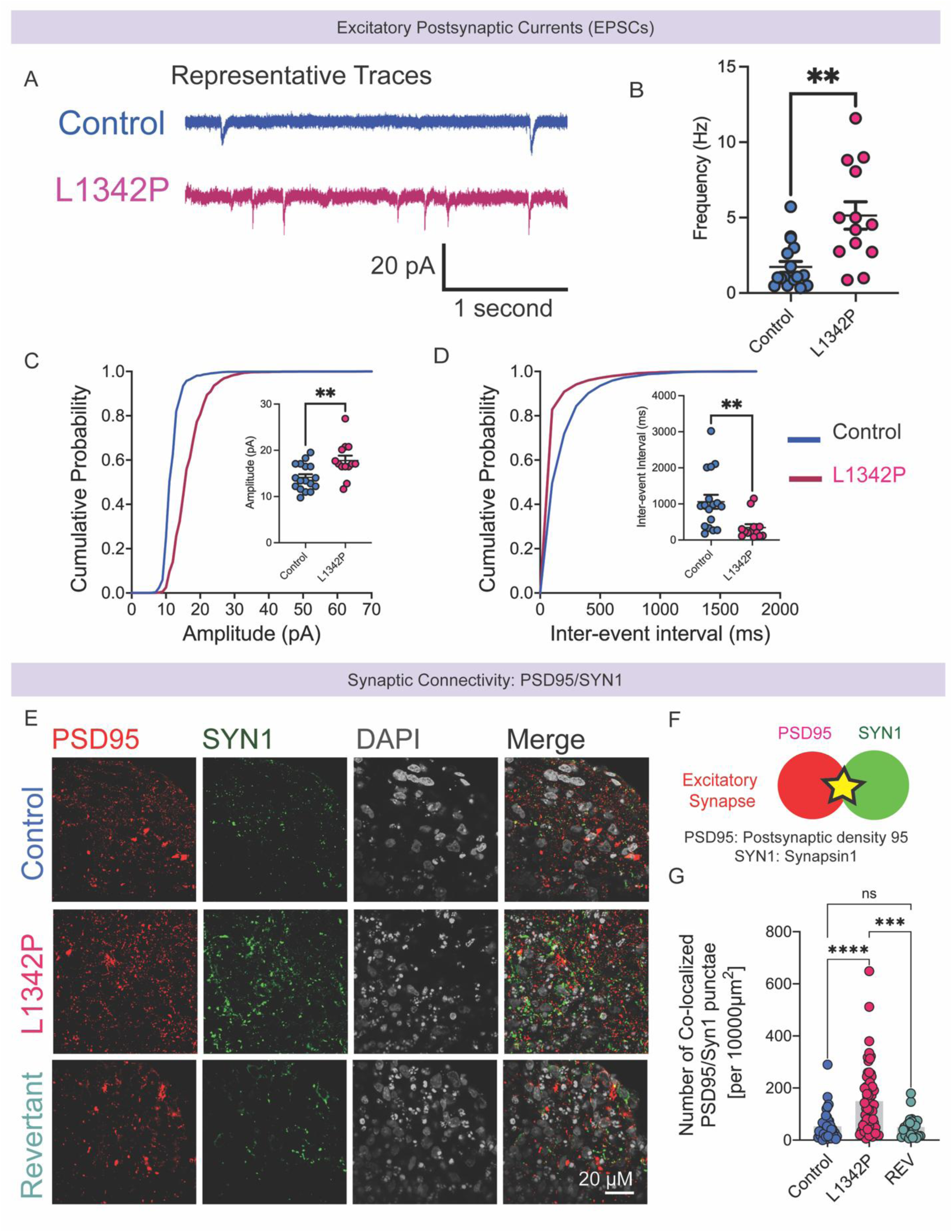
Nav1.2-L1342P cortical organoid neurons display enhanced excitatory synaptic transmission and enhanced synapse formation compared to isogenic controls. **(A)** Representative excitatory post-synaptic currents (EPSCs) traces were recorded from Control and Nav1.2-L1342P cortical organoid neurons. **(B)** EPSC frequency is increased in Nav1.2-L1342P cortical organoid neurons. **(C)** The cumulative probability distribution of EPSC amplitude, with quantification indicating an increase in amplitude within Nav1.2-L1342P cortical organoids. **(D)** The cumulative probability distribution of EPSC inter-event interval, with quantification revealing decreased inter-event intervals in Nav1.2-L1342P cortical organoids. **(E)** Increased synapse formation in cortical organoids carrying the Nav1.2-L1342P mutation at Day 120, as evidenced by the markers Post-synaptic Density Protein-95 (PSD95) in magenta and Synapsin1 in green. **(F)** Components of an excitatory synapse. **(G)** The number of co-localized punctae (synapses) was increased in the Nav1.2-L1342P Cortical organoids compared to controls and revertant lines. Data are reported as mean ± error (S.E.M). *One-Way ANOVA*, * p < 0.05; ** p < 0.01; ****p < 0.001.

Increased sEPSC might arise as a result of increased formation of excitatory synapses, or enhanced spontaneous neuronal firing. Thus, we performed immunocytochemistry of our cortical organoids **(Figure 2E)** against two vital excitatory synaptic proteins: Synapsin −1 (SYN1) and post-synaptic density protein (PSD95) **(Figure 2F)**. SYN1 is a presynaptic phosphoprotein crucial for synaptogenesis and synaptic plasticity (Parenti et al., 2022). PSD95 is a scaffolding protein located at excitatory post-synapses, involved in the stabilization, recruitment, and trafficking of N-methyl-D-aspartic acid receptors (NMDARs) and α-amino-3-hydroxy-5-methyl-4-isox-azoleproprionic acid receptors (AMPARs) to the post-synaptic terminals(Coley & Gao, 2019). We quantified the number of colocalizations (synapses) among conditions normalized by the area occupied by a MAP2 stain, which delineates the start and end of the cortical organoid slice. We found that the colocalization density of SYN1^+^PSD95^+^ synaptic vesicle proteins was significantly increased in Nav1.2-L1342P cortical organoids compared to control and revertant organoids in D120, suggesting an increase in the formation of excitatory synapses in Nav1.2-L1342P organoid neurons (Control: 52.89 ± 9.438, n = 37 images, 6 organoids, 2 clones across 3 differentiations; L1342P: 149.5 ±16.22, n = 61 images, 10 organoids, 2 clones, across 3 differentiations; REV: 50.15 ± 9.146, n = 23 images, 5 organoids, 1 clones, across 2 differentiations, **Figure 2G)**. Our data indicate that the Nav1.2-L1342P mutation affects synapse formation by enhancing pre- and post-synaptic colocalization, which is consistent with our EPSC data indicating a functional alteration in synaptic function.

### Cortical Organoids Carrying the Nav1.2-L1342P Mutation Display Marked Network Hyperexcitability

Single-cell patch-clamp recordings revealed elevated evoked action potential (AP) firing, intrinsic excitability, and spontaneous excitatory post-synaptic currents (sEPSCs) in individual cortical organoid neurons carrying the Nav1.2-L1342P mutation. To directly test whether these changes contribute to network excitability, we performed multi-electrode array (MEA) recording of our cortical organoids. This higher-throughput extracellular recording assay enables the simultaneous recording of spike firing and field potentials across an entire neuronal population (Hiranuma et al., 2024), providing an avenue to assess overall neuronal network excitability **(Figure 3A)**. For these experiments, we plated three conditions: WT control, Nav1.2-L1342P, and revertant P1342L cortical organoids directly on top of the electrodes for the recordings **(Figure 3B)**.

**Figure 3.**
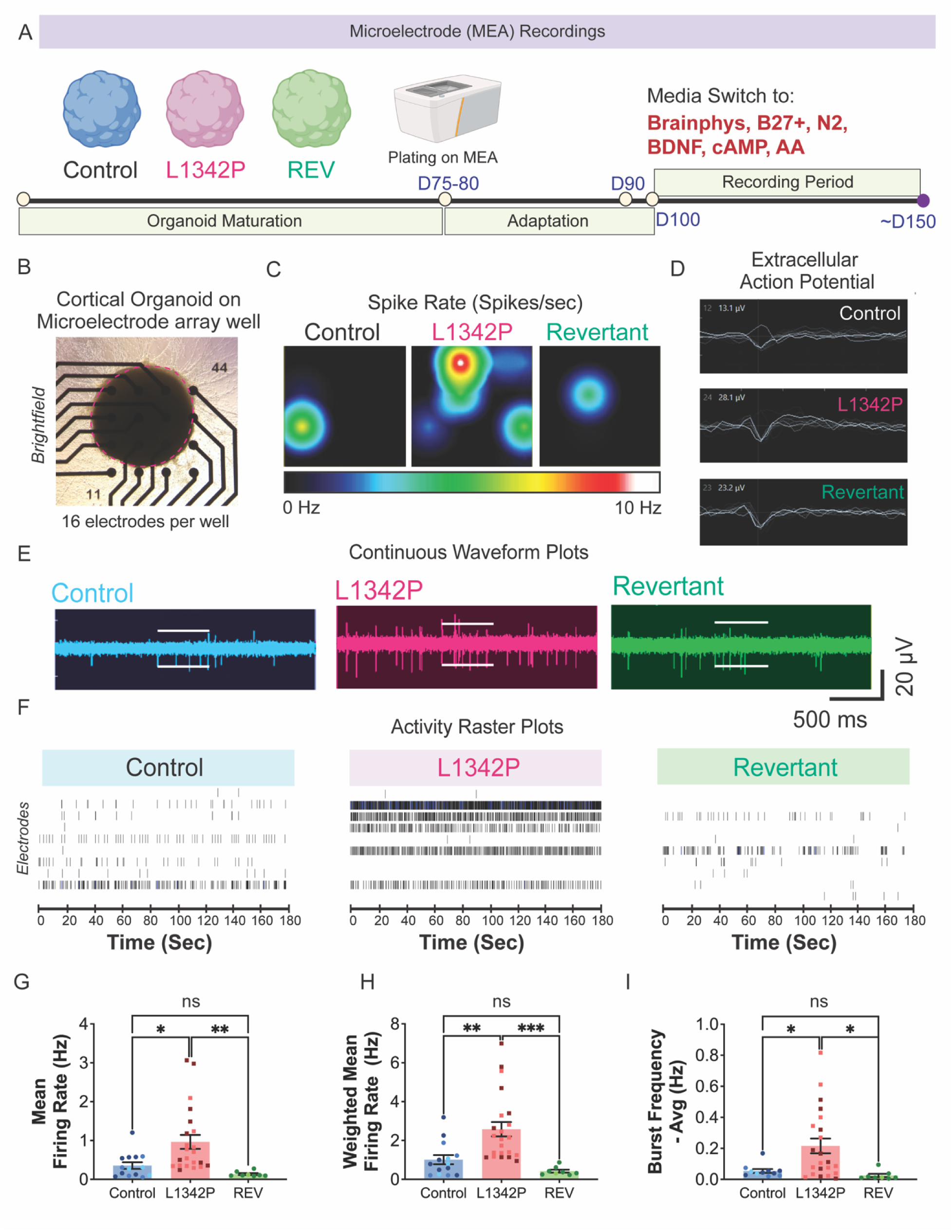
MEA recording reveal that cortical Organoids carrying the Nav1.2-L1342P mutation display marked network hyperexcitability. **(A)** Description of organoids used in the study. **(B)** Brightfield image of an organoid placed directly on top of MEA electrodes. **(C)** The firing of a population of neurons can be visualized in a heatmap view. Each colored circle represented an active electrode, with blue indicating a low firing frequency and yellow/red indicating a high firing frequency. Nav1.2-L1342P cortical organoids had increased spike rate values compared to controls and revertants. **(D)** Raw spikes. Each spike indicated a spontaneous activity. Nav1.2-L1342P cortical organoids had increased spontaneous activity values compared to controls and revertants. **(E)** Representative spike raster plots. Nav1.2-L1342P cortical organoids had increased activity compared to controls and revertants. **(F)** The mean firing rate was increased in Nav1.2-L1342P cortical organoids compared to controls and revertants. **(G)** The weighted mean firing rate was increased in Nav1.2-L1342P cortical organoids compared to controls and revertants. **(H)** The burst frequency of Nav1.2-L1342P cortical organoids was increased compared to controls and revertants. Each dot represents an organoid. Data was reported as mean ± error (S.E.M). *One-Way ANOVA* was performed with * p < 0.05; ** p < 0.01; ****p < 0.001.

We demonstrated that Nav1.2-L1342P cortical organoids exhibited significantly higher neuronal activity compared to controls and the revertant line, as illustrated in heatmap views **(Figure 3C)**, extracellular action potential traces **(Figure 3D)**, continuous waveform plots **(Figure 3E),** and activity raster plots **(Figure 3F)**. Quantitatively, MEA recordings revealed that the Mean Firing Rate (MFR) of Nav1.2-L1342P cortical organoids was substantially elevated compared to controls and revertant cultures (Control: 0.3553 ± 0.08487 Hz, *n* = 14 organoids, three clones across six differentiations; L1342P: 0.9646 ± 0.1793 Hz, *n* = 22 organoids, two clones, across six differentiations; REV: 0.1362 ± 0.02374 Hz, *n* = 8 organoids, two clones, across two differentiations, *One-Way ANOVA,* **Figure 3G)**, and the same trend was evident in the Weighted Mean Firing Frequency (WMFF) (Control: 1.013 ± 0.2363 Hz, *n* = 14 organoids, three clones across six differentiations; L1342P: 2.579 ± 0.3687 Hz, *n* = 22 organoids, two clones, across six differentiations; REV: 0.4244 ± 0.07684 Hz, *n* = 8 organoids, two clones, across two differentiations, *One-Way ANOVA,* **Figure 3H)**. Bursting frequency was further quantified, as it is a seizure-related firing characteristic of neuronal cultures. The average bursting frequency was significantly increased in Nav1.2-L1342P cultures compared with controls and the revertant line (Control: 0.05412 ± 0.01241 Hz, *n* = 11 organoids, three clones across six differentiations; L1342P: 0.2157 ± 0.04780 Hz, *n* = 22 organoids, two clones, across six differentiations; REV: 0.02502 ± 0.01051 Hz, *n* = 8 organoids, two clones, across two differentiations, *One-Way ANOVA,* **Figure 3I)**. These findings suggest that the Nav1.2-L1342P cortical organoids enhance network excitability in the 3D cortical organoids.

### RNA-Sequencing Analysis of hiPSC-Derived Cortical Organoids Reveals Impairments in Synaptic, Glutamate, Development-Related Pathways, and Apoptosis

Following the functional characterization of Nav1.2-L1342P cortical organoids, which revealed profound hyperexcitability, we sought to explore the molecular mechanisms underlying this enhanced neuronal excitability. We employed RNA sequencing to assess transcriptomic signatures and gene regulatory networks associated with the Nav1.2-L1342P mutant organoids. We collected six samples per condition from at least two differentiations **(Figure 4A)**. The sample Sequence Similarity Matrix **(Figure 4B)** indicates that LP and Control organoid samples form two distinct clusters. The 3D PCA Plot also shows a clear separation between mutant (pink) and control (blue) samples along PC1, PC2, and PC3 **(Figure 4C)**. Our analysis identified a total of 5701 globally differentially expressed genes (DEGs) with a False Discovery Rate (FDR) < 0.05. We found 2816 genes were downregulated, and 2354 were upregulated **(Figure 4D)**. The top 40 DEGs comprised 10 downregulated genes and 30 upregulated genes (**Figure 4E**).

**Figure 4.**
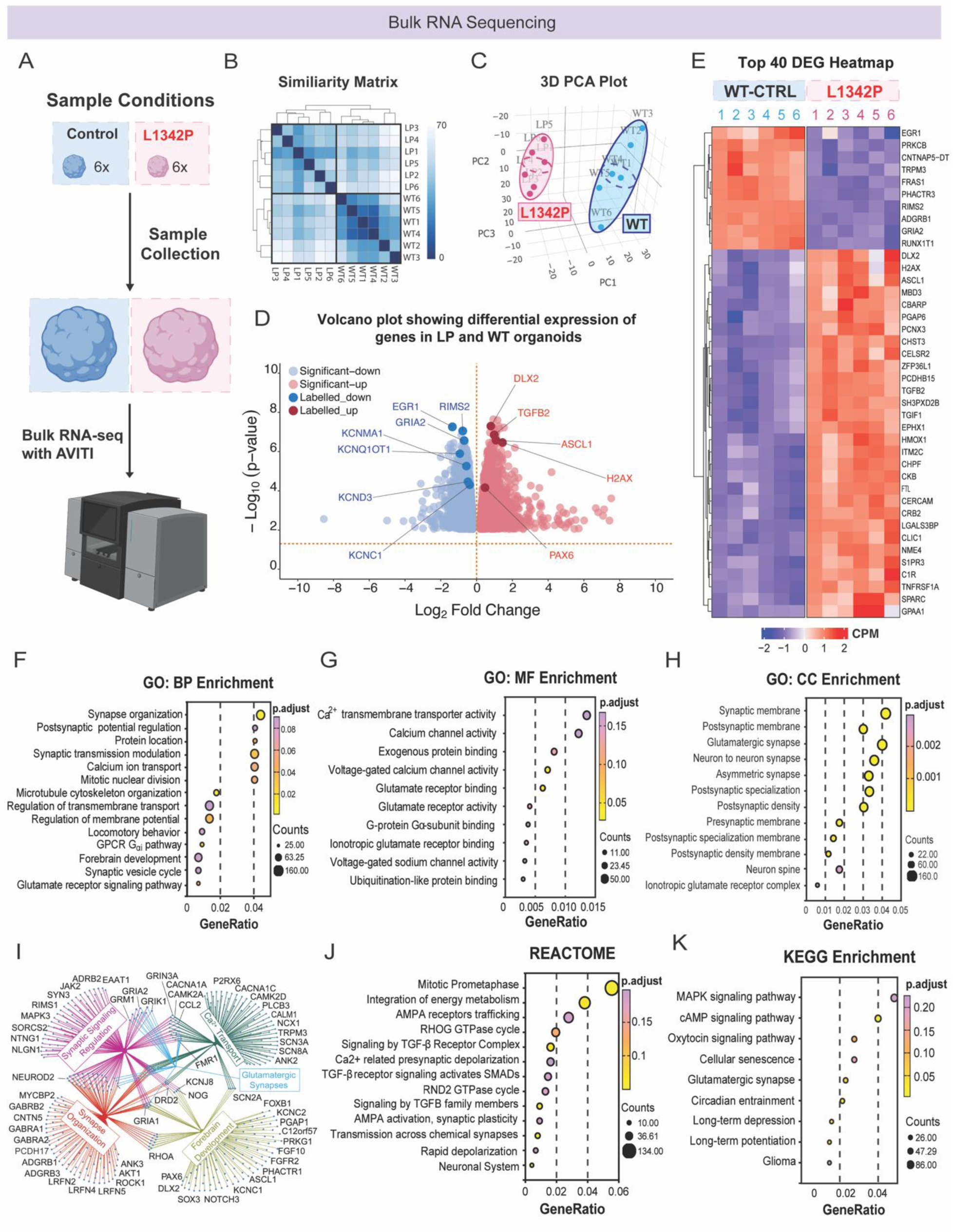
RNA-sequencing analysis of hiPSC-derived cortical organoids reveal impairments in synaptic, glutamate, and forebrain developmental pathways. **(A)** Bulk RNA-seq experimental conditions. Control organoids (n = 6 organoids, three differentiations) and L1342P organoids (n = 6 organoids, two differentiations) were sequenced using the AVITI system. **(B)** Hierarchical clustering of cortical organoid sample replicates. Darker blue indicates higher similarity. **(C)** Principal Component Analysis (PCA) plot illustrating the clustering of replicates based on their variance in two dimensions. **(D)** DEG Heatmap showing the top 50 down (blue) and upregulated (red) genes across conditions. **(E)** Volcano plot highlighting significantly differentially expressed genes (DEGs, FDR < 0.05) in Nav1.2-L1342P hiPSC-derived cortical organoids compared to controls. Notable downregulated genes include SCN2A and potassium channel-related genes. In contrast, upregulated genes include: DLX2 (Distal-Less Homeobox 2), H2AX (histone family member X), PAX6 (Paired Box 6), and ASCL1 (Achaete-scute family bHLH transcription factor 1), which are important in neurodevelopment, neuronal function, and cellular function. **(F)** Gene Ontology (GO) analysis of biological processes reveals changes in synapse organization, microtubule-cytoskeleton arrangements, glutamate receptor signaling, and forebrain development (top). **(G)** GO molecular function analysis reveals enrichment in ion channel activity and alterations in the glutamatergic pathway. **(H)** GO cellular component identifies synaptic pathway alterations in the Nav1.2-L1342P cortical organoids. **(I)** Network analysis reveals numerous globally differentially expressed genes (DEGs) involved in synaptic signal regulation (magenta), calcium transport (dark green), synapse organization (orange), and forebrain development (lime). The colored edges connecting genes represent functional relationships or interactions within each cluster, highlighting how these DEGs collectively contribute to altered neuronal physiology. **(J)** REACTOME pathway analysis reveals disruptions in cell division, metabolism, AMPA (α-amino-3-hydroxy-5-methyl-4-isoxazolepropionic acid) receptor trafficking, and neuronal synapses. **(K)** KEGG pathway enrichment analysis reveals alterations in MAPK and cAMP signaling pathways, as well as glutamatergic synapses. The circle size is represented by the number of DEGs highly ranked by p.adjust values.

To investigate the biological processes affected by the Nav1.2-L1342P organoids, we conducted a functional enrichment analysis of our differentially expressed genes using the clusterProfiler **(Figure 4F-K)**. Gene Ontology (GO) Biological Processes (GO.BP) analysis revealed disruptions in pathways related to synapse organization, microtubule-cytoskeleton dynamics, glutamate receptor signaling, cell division, and forebrain development **(Figure 4F)**. Notably, several genes associated with forebrain development were identified, including DLX2, ASCL1, H2AX, SCN2A, POU3F2, CHD5, SLC4A10, and PAX6 **(Figure 4D, E, I)**, further suggesting the role of Na_V_1.2 in development. Many of these developmental genes were increased, PAX6 (logFC: 0.4718, p value: 6.876e-05), a marker of neural progenitor cells; H2AX (logFC: 1.446, p value: 3.425e-07), a key regulator of mitotic prometaphase; and ASCL1 (logFC: 1.082, p value: 2.631e-07), a master regulator of neurogenesis. Interestingly, potassium channel-related genes, e.g., (KCNC1, KCNC2) were differentially downregulated within this GO network. Given that these organoids were sampled at a developmental stage when mature neuronal markers are apparent (130-167 days), this pattern may suggest delayed or impaired neuronal maturation in the L1342P condition.

GO Molecular Function (GO.MF) analysis revealed perturbations in calcium and sodium ion channel activity, as well as glutamatergic signaling, indicating dysregulation of cellular excitability, which is consistent with our previous findings on hyperexcitability and EPSC-related mechanisms **(Figure 4G)**. Moreover, our GO Cellular Component (GO.CC) analysis, which describes subcellular structures and macromolecular complexes (Roncaglia et al., 2013), also identified significant alterations in synaptic pathways **(Figure 4H)**, consistent with our SYN1/PSD95 results. Network pathway analysis of differentially expressed genes further grouped them into four main functional clusters, each represented by a distinct color and connected by nodes representing interactions with each other. Synaptic signaling regulation genes (magenta), such as GRIA2, GRIN3 and JAK1 modulate synaptic activity and neurotransmission. Calcium transport genes (dark green), including CAMK2A, and CACNA1A, regulatate calcium signaling. Synapse organization genes (orange), such as ANK3, GABRA2 and GRIA1, contribute to the structural organization and maintenance of neural connections. Forebrain development genes (lime) include key transcription factors and developmental regulators such as SOX3, PAX6, NOTCH3 which are important in cortical development **(Figure 4I)**.

REACTOME pathway analysis indicated disruptions in cell division, metabolism, AMPA receptor trafficking, and neuronal synapses **(Figure 4J)**. Interestingly, KEGG pathway analysis revealed dysregulation in MAPK and cAMP signaling, as well as glutamatergic synapses and cellular senescence **(Figure 4K).** In our transcriptomics analysis, the KEGG senescence pathway was found to be significantly enriched in L1342P organoids, with the upregulation of many genes associated with cellular senescence and autophagy, for example, H2AX (logFC: 1.446, p value: 3.425e-07), a double-stranded DNA marker; NME4, a nucleoside diphosphate kinase (logFC: 0.918, *p value*: 5.6266e-08 too many digits), was among the top 10 upregulated genes and plays a role in the early stages of mitophagy and autophagy (Lacombe et al., 2018). GPX1, one of the top 100 DEGs, was also upregulated and encodes a glutathione peroxidase that scavenges reactive oxygen species during oxidative stress, particularly in status epilepticus (Kim et al., 2024). Moreover, several apoptosis-related genes were significantly increased, including CASP1 (logFC: 1.430; *p value*: 0.00014), an early apoptotic marker, as well as pro-apoptotic BCL-2 family members (BAX/BAK). Several neuroimmune-related genes linked to autophagy and phagocytosis in epilepsy were also prominently upregulated, including CXCL3 (logFC: 1.876, p value: 0.0023), CCL2 (logFC: 2.271, p value: 0.0021), and C1R (logFC: 1.167, p value: 2.483E-08). Taken together, these findings suggest that L1342P organoids may be undergoing significant cellular stress and early pro-apoptotic, and neuroimmune signaling, possibly relating to the reported human cerebral atrophy phenotypes. Together, these findings suggest the profound impact of the Nav1.2-L1342P mutation on neuronal development and function, further supporting its role in neurodevelopmental pathophysiology.

## Discussion

This study demonstrates that the recurring Nav1.2-L1342P mutation has a profound impact on neuronal excitability and the expression of neurodevelopmental genes. In this 3D system, we observed a hyperexcitability phenotype characterized by enhanced repetitive neuronal action potential firing, increased intrinsic excitability, increased network neuronal firing, and excitatory post-synaptic currents (EPSCs). Immunocytochemical analyses revealed an increased number of glutamatergic synapses, offering a cellular correlate for the observed network activity. Along with increased synapses, RNA sequencing of Nav1.2-L1342P cortical organoids revealed significant dysregulation of genes associated with synaptic signaling, glutamatergic neurotransmission, neurodevelopment, and cellular apoptosis/senescence, further supporting the role of the Nav1.2-L1342P mutation in affecting neuron development in both 2D and 3D models. The use of a CRISPR/Cas9 gene-corrected P1342L revertant control further strengthens our conclusions, by ensuring that observed phenotypic differences are specifically attributable to the mutation, thereby enhancing the rigor of our study.

SCN2A-L1342P affects multiple patients including 5 reported in the literature (Crawford et al., 2021; Hackenberg et al., 2014; Matalon et al., 2014; Wolff et al., 2017). Recently, a sixth male patient, born in 2024, was identified with the L1342P variant. This patient developed seizures at five months of age, with a BASED (Burden of Amplitudes and Epileptiform Discharges) score escalating from 1 to 4 within five months. Comorbidities included sleep disorders, dystonia, chorea, hypotonia, gastrointestinal issues, and sensory hypersensitivity. Despite treatment with multiple anticonvulsants, including Prednisolone, Lacosamide, Clonazepam, Clobazam, Vigabatrin, Epidiolex, and adrenocorticotropic hormone (ACTH), seizure control remains poor. The recurrent nature of the L1342P variant suggests that this site may be a mutation hotspot. Moreover, because most patients with L1342P do not achieve sufficient seizure control, understanding the molecular and cellular phenotypes associated with this mutation may help inform the development of targeted therapeutic interventions.

Cortical organoid models have been widely used to study the relationships between cellular changes and neuronal function in disease. In a forebrain organoid model of Fragile X syndrome, the loss of FMRP disrupts cortical neuron development and enhances neuronal excitability, as evidenced by an increase in synaptic puncta density, and leads to widespread alterations in gene expression (Kang et al., 2021). In our study, we observed similar features in cortical organoids carrying the Nav1.2-L1342P variant, including an increase in co-localized synaptic puncta, neuronal hyperexcitability, and global transcriptional changes resulting from a single amino acid substitution. However, this pattern is not consistent across all *in vitro* epilepsy models. In an organoid model of Focal Cortical Dysplasia (FCD) type II, one of the most common causes of drug-resistant epilepsies. Organoids exhibit impaired cell proliferation, dysmorphic neurons, spontaneous hyperexcitability, and enhanced network connectivity, which were paradoxically accompanied by reduced actin cytoskeleton polarization and fewer synaptic puncta (Avansini et al., 2022). Together, these studies suggest that the connection between electrical activity and its underlying structural correlates is highly complex and that structure-function relationships can be challenging to predict. They also emphasize the value of 3D brain organoids as a powerful platform to model complex case-specific neurological phenomena and investigate epilepsy risk variants that disrupt both cellular and circuit connectivity.

Transcriptomic profiling of Nav1.2-L1342P cortical organoids revealed significant alterations in neurodevelopmental gene expression, glutamatergic signaling, excitotoxicity-related pathways, and widespread downregulation of multiple potassium channel-related genes. Similar transcriptomics changes have been reported in other SCN2A *in vitro* and *in vivo* models. For instance, analysis of hyperexcitable 2D iPSC-derived neuronal models carrying SCN2A-R1882Q and SCN2A-E1803G variants also revealed upregulation of neurodevelopmental genes such as NEUROD6 and FOXG1. Gene-ontology network analysis in these models further identified dysregulation in extracellular matrix organization, regulation of cell proliferation, and adhesion (Asadollahi et al., 2023), implicating broader defects in cellular development across SCN2A-related pathologies. In our L1342P organoid model, KEGG pathway analysis revealed significant enrichment of senescence and autophagy-related processes. Notably, we observed increased expression of caspase-associated autophagy genes, including CASP1, as well as downstream BCL2 effector genes BAX and BAK. A similar caspase-related gene expression profile was observed in the R1882Q 2D neuron model, which revealed upregulation of NLRP2, a key component of the innate immune system that activates caspase-1 and promotes apoptosis (Mao et al., 2024). Additionally, the elevated expression of oxidative stress-related genes NME4 and GPX1 in our L1342P organoids aligns with findings from the SCN2A-E1803G model, which also exhibited activation of senescence and NRF2-mediated oxidative stress response pathways (Asadollahi et al., 2023). Abnormal neurotransmitter dysregulation, particularly involving glutamate, has also been reported in the context of sodium channel dysfunction. In our previous work, we observed increased glutamate release in hiPSC-derived neuronal monolayers carrying the L1342P variant (Que et al., 2025), further implicating glutamatergic signaling in excitotoxicity-related disease pathology. In this current study, cortical organoids carrying the L1342P mutation exhibited downregulation of multiple potassium channel-related genes, including several voltage-gated (KCNC1, KCNA4, KCNH5), inward-rectifying channels (KCNJ3, KCNJ9), and calcium-gated potassium channels (KCNMB2, KCNN3, KCNU1), among others. These channels regulate the flow of potassium ions through the cell, influencing membrane excitability, and are recognized as major mediators of neuronal excitability. Reduced expression of the K^+^ channels may contribute to neuronal hyperexcitability by impairing the repolarization of the action potential, however how the complex interplay between all these channels in the context of SCN2A-L1342P needs to be explored in more detail to understand K+ channel contributions to excitability. Genetic interactions between SCN2A and potassium channels, such as KCNQ2 (Kv7.2) (Kearney et al., 2006) and KCNA1 (Kv1.1)(Spratt et al., 2021; Zhang et al., 2021), have been described in mouse models of SCN2A and are also reflected in our study.

Our study has limitations that should be acknowledged. Cortical organoids are heterogeneous at the cellular level, comprising neural progenitor cells and neurons, as well as glial progenitor cells that give rise to astrocytes and oligodendrocytes (Yoon et al., 2019). Transcriptomics at the bulk RNA sequencing level cannot provide detailed cell-type-specific information; therefore, it is not possible to assign cell type identity to the specific gene expression and pathway analysis we performed. One study has shown that SCN2A deletion impairs the differentiation of a subpopulation of oligodendrocyte progenitor cells during late development/differentiation stages, but not during proliferation or production of these progenitor cells (Gould & Kim, 2021). Single-cell RNA sequencing could help address this limitation in future studies.

While the rescue of the hyperexcitability phenotype was not explored in this current study, future studies could leverage this platform to evaluate therapeutic interventions, ranging from pharmacological treatments to gene therapy approaches, aimed at reducing and modulating neuronal hyperexcitability. When applied to *in vitro* models, various pharmacological and gene replacement interventions have been shown to rescue the abnormal hyperexcitability phenotypes associated with SCN2A (Chen et al., 2024; Que et al., 2021). Additionally, antisense oligonucleotides (ASOs) are being developed to target pathogenic SCN2A GoF variants implicated in early-onset developmental and epileptic encephalopathy. For instance, intrathecal elsunersen, used to treat a female preterm infant with early-onset *disease*, resulted in a >60% reduction in seizure frequency during follow-up until the age of 22 months (Wagner et al., 2025). Utilizing a brain organoid model, it might be possible to test personalized ASO *in vitro* first before patient application.

In summary, we have established a preclinical *in vitro* iPSC-derived cortical neuron model to investigate the effects of the L1342P mutation. This GoF SCN2A epilepsy-associated mutation causes developmental and epileptic encephalopathy and abnormal brain structure in human patients. Our findings validate the utility of hiPSC-derived technologies for studying *SCN2A*-related pathology, proving that the gain-of-function L1342P variant influences complex molecular and cellular mechanisms beyond abnormal hyperexcitability. This highlights the multifaceted impact of SCN2A variants and their capacity to disrupt cellular processes and lead to disease. This study represents an advancement in using cortical organoids to model GoF SCN2A mutations and provides a platform for testing novel interventions in human cell-based preclinical models.

## Acknowledgments

This work was supported by National Institutes of Health National Institute of Neurological Disorders and Stroke (NINDS) Grants R01 NS117585 and R01 NS123154 (to Y.Y.). M.I.O.A. is supported by the Fulbright-Colciencias Scholarship Program. We thank Dr. Mark Estacion for the advice on calcium imaging assays. We also thank the support from the Purdue University Institute for Drug Discovery (PIDD), the Purdue Institute for Integrative Neuroscience (PIIN), and the Borch Department of Medicinal Chemistry and Molecular Pharmacology. The Yang Lab thanks the *FamilieSCN2A* foundation for the Action Potential and Hodgkin-Huxley Award.

## Author contributions

M.I.O.A. and Y.Y. designed research; W.C.S. contributed unpublished reagents/analytic tools. M.I.O.A, M.R, Z.Q, Z.Z, H.H, V.S, S.H, M.W, M.O, H.K, M.H, C.D, T.N, K.W, B.Z, L.Y, N.C, X.C, J.Z, J.W, B.D, Y.Z performed research; M.I.O.A, M.R, V.S, M.W, M.O, H.H, T.N, C.Y, H.K, F.G and Y.Y analyzed data; M.I.O.A, M.R., and Y.Y wrote the paper.

## Methods

### Generation of Cortical Organoids from HiPSCs

To model the pathophysiology of the monogenic Nav1.2-L1342P mutation, we utilized CRISPR/Cas9-edited hiPSCs derived from a KOLF2-C1 male donor reference line. Three lines sharing the same genetic background were generated: Controls (Wild-type), CRISPR/Cas9-engineered Nav1.2-L1342P (Que et al., 2021), and a P1342L revertant line. The revertant line was used to confirm the functional effects of the specific Nav1.2-L1342P engineered mutations on the resulting disease traits. This approach ensured that the observed phenomena were not a result of genetic background effects, but rather from the single-amino acid substitution, further facilitating genotype-phenotype correlations. Quality controls were performed to ensure line integrity, including Sanger sequencing, karyotyping, and immunocytochemistry for hiPSC pluripotency. Undifferentiated hiPSC colonies from controls and Nav1.2-L1342P lines displayed normal and homogenous morphology with defined edges and low levels of spontaneous differentiation. They consistently expressed pluripotency markers, including OCT4, SSEA4, and TRA-1-60, making them suitable for cortical neuron monolayer differentiations.

Cortical organoids were prepared using the protocol developed by the Pașca lab at Stanford(Miura et al., 2022). For all experiments, hiPSC colonies were maintained daily in StemFlex Medium (Gibco, Catalog No. A334940) and passaged manually in clumps with Versene every 3-4 days. To generate cortical organoids, hiPSCs were aggregated at a cell density of 100 cells/μL in Essential 8 medium (Gibco, Catalog No. A1517001) to form organoid embryoid bodies (EBs). Round-bottom ultra-low-attachment plates (Corning Costar, Catalog No. CLS7007) and centrifugation at 100G for 3 minutes facilitated the formation of EBs overnight. The EBs were maintained in Essential six medium (Gibco, Catalog No. A1516401) for the first six days, supplemented with activin/nodal/TGF-β and BMP pathway inhibitors 2.5 µM Dorsomorphin (DM), 10 µM SB-4321542, and 1.25 μM XAV-939 for neuronal induction via DUAL-SMADi method. EBs were harvested and transferred to flat-bottomed six well-formatted suspension culture plates (Corning, Catalog No. 347) and maintained in Neurobasal -A medium with Glutamax (Gibco, Catalog No. 35050061) and PenStrep (10000 U/mL) prophylactic (Gibco, Catalog No. 15140163), 1X B27 minus A supplement (Gibco, Catalog No. 12587010) with 20 μg/mL Human Recombinant FGF2 (Fibroblast growth factor 2) and 20 μg/mL EGF (Epidermal growth factor) with complete medium changes every day for 17 days followed by every two days until day 23. Afterward, the added medium supplements were replaced with 20 ng/mL BDNF (Brained Derived Neurotrophic Factor), 20 ng/ mL NT3 (Neurotrophin-3), 50 μM cAMP (Cyclic adenosine monophosphate), 10 μM DHA (cis-4,7,10,13,16,19-Docosahexaenoic acid) and 200 μM AA (L-ascorbic acid (AA) 2-phosphate trisodium salt) and changed every two days to achieve organoid differentiation until day 46. After that, the organoids were maintained in Neurobasal-A basal medium supplemented with 1X B27+ and without growth factors until day 150, with medium changes every 4-5 days.

### Viral Labelling of Cortical Organoids

We gathered three ∼75-day-old organoids in a 1.5 mL centrifuge tube containing 200 μL of organoid maturation medium, which consisted of neurobasal medium supplemented with B27+. We added 1μL of pAAV-CaMKIIa-EGFP virus at a 1×10¹³ vg/mL titer to label cortical excitatory organoid neurons for electrophysiology experiments. We incubated them overnight at 37 °C, 5% CO_2_. The next day, we added an extra 800 μL of organoid maturation medium to the mix and further incubated for 24 hours (Sloan et al., 2018). After this incubation period, cortical organoids were collected and then plated on PLO-Laminin-coated coverslips and were maintained until visible neurites emanated from the plated organoid core and reached maturity (e.g., 120-160 days).

### Electrophysiology of hiPSC-Derived Cortical Organoid Neurons

To examine single neuronal activity patterns, 75-day-old organoids were plated on glass coverslips (Neuvitro, Catalog No. 12-GG-PRE) pre-coated with poly-L-ornithine (PLO) and laminin. Briefly, coverslips were coated overnight at room temperature with a 1:5 dilution of 0.01% (PLO; Sigma-Aldrich, Catalog No. P4957) in phosphate-buffered saline (PBS). The next day, PLO-coated plates were incubated with 10 μg/mL mouse laminin (Corning, Catalog No. 354232) in DMEM/F12 for 2 hours at 37°C (Que et al., 2025). Organoids were placed at the center of each coverslip in 20 μL of Neurobasal-A medium supplemented with B27+ and allowed for up to 4 hours in a 37°C incubator. Following this, an additional 40 μL of medium was gently added to each well to support continued attachment overnight. Sterile water was added to the MEA chambers to maintain a moist environment and prevent evaporation. The next day, 300 μL of medium was added to the coverslips and left undisturbed for several days, until visible neurites could be seen emanating from the organoid core and fusing into the coverslip-coated matrix, generating a monolayer of neurons. Cortical organoids were then maintained in electrophysiology-supporting medium containing Brainphys (Stemcell technologies, Catalog No. 05790), 1X B27+, 1X N-2 Supplement (Gibco, Catalog No. 17502048), 20 ng/mL BDNF, 50 μM cAMP, and 200 μm AA with half medium changes every 3-4 days.

To select neurons for patch -clamp experiments, we focused on patching the pAAV-CaMKIIa-EGFP-labeled excitatory cell populations that protruded from the replated organoid core. The procedure for electrophysiology is as follows: Whole-cell current-clamp studies were performed on GFP-labeled neurons carrying either Control or Nav1.2-L1342P mutant channels. Signals were amplified with a MultiClamp 700B amplifier (Molecular Devices) with a 2 kHz filter and digitized at 10 kHz with a Digidata 1550B A/D converter (Molecular Devices). Data were collected using Clampex 11.2 software (Molecular Devices) and stored on a PC for future analysis. We obtained several relevant parameters, including voltage threshold, amplitude, spike width, resting membrane potential, rheobase, and EPSC traces. Whole-cell voltage-clamp recordings obtained the EPSCs of different experimental groups.

### Cortical Organoid Microelectrode Array (MEA) Recordings

75-day-old organoids were plated into a 48-well CytoViewMEA plate (Catalog No. M768-tMEA-48B) obtained from Axion BioSystems to assess organoid network activity. The MEA plate wells were coated with the same PLO-Laminin composition described previously. Cortical organoids were carefully placed directly on the electrodes and maintained for the first week (7 days) in neuronal maturation medium composed of Neurobasal-A and B27+ supplement. After that initial period, medium was replaced with BrainPhys medium supplemented with B27+ and N-2 supplements, BDNF, AA, and cAMP in the same concentrations as previously indicated. All recordings were performed in a time window spanning 120-140 days. Before all recordings took place, MEA plates were stabilized in the machine unit for at least 5 minutes under controlled conditions of 37 °C and 5% CO_2,_ which are suitable for live cell recordings. For analysis, we used recordings that were 3 minutes long. The threshold for determining voltage spikes was set to 5.5 standard deviations above the background noise. Active electrodes were defined as more than five spikes per minute (0.0833 Hz) of recording, and wells with <25% active electrodes were not used for processing. Parameters measured included the mean firing rate, weighted mean firing weight and bursting frequency and were obtained using the Neural Metrics Tool, version 2.4. from Axion Biosystems (Que et al., 2021).

### Preparation and Immunostaining of Cortical Organoid Slices

For immunostaining experiments, organoids were collected at the following time points: day 30 (D30) and day 120 (D120). Using a wide-bore pipette tip (Axygen, Catalog No. T205WBCRS) or a cut 1 mL tip, organoids were retrieved from their plates, placed on a 1.5 mL centrifuge tube, medium removed, and 1 mL of 4% Paraformaldehyde (PFA) was added for overnight fixation at 4 °C. The following day, PFA was removed, and organoids were dehydrated in a 30% sucrose solution in Dulbecco’s phosphate-buffered saline (DPBS) to avoid crystal formation during freezing. When adding the sucrose at this point, organoids should be floating on the surface. After 3-4 days, organoid samples are considered ready when they have sunk to the bottom of the tube. Between 2-5, organoids are placed on molds and covered in Sakura tissue-tek O.C.T. Compound (Sakura Finetek USA, Catalog No. 4583) and left on regular ice to allow the organoid to settle at the bottom. Afterward, samples were placed on top of dry ice for 10 minutes and stored at – 80 °C until needed.

Organoid slices were obtained using a Leica Cryostat, set to a temperature ranging from −18 to −20 °C. The cryostat was configured to collect slices at 20 μm thickness for PSD95/SYN1 quantification and 40 μm thickness for cortical rosettes and neurons. Slices were obtained without the glass insert to prevent tissue fracturing and were collected using a glass slide (VWR MicroSlides, Catalog No. 48311-703).

For immunostaining, samples are air-dried and washed three times with DPBS for at least 5 minutes to dissolve the remaining OCT embedding solution and rehydrate the sample. After drying, the slices were circled with a hydrophobic marker. For the blocking step, a solution of 0.3% Triton X and 5% BSA in PBS was added to the slices for 1 hour at room temperature. The sections were then incubated overnight in a humidified chamber at 4 °C with primary antibodies in a blocking solution (0.1% Triton X, 1% BSA in PBS). Primary antibodies included: chicken anti-MAP2 (Microtubule Associated Protein-2, Novus biologicals, Catalog No. NB300-213, 1:1000), rabbit-anti SOX2 (SRY-Box Transcription Factor 2 SRY-Box Transcription Factor 2, Cell Signaling Technology, Catalog No.D6D9, 1:500), mouse anti-Ki67 (Human Ki67 Pure B56, BD Biosciences, Catalog No. 556003, 1:200), mouse anti-NEUN (NeuN, Catalog No. E4M5P, 1:500), mouse anti-Synapsin1 (SYN1, Synaptic Systems, Catalog No. 106011, 1:200) and rabbit anti-PSD95 (Invitrogen, Catalog No. PIPA585749, 1:200). The next day, slides were carefully washed with PBS at RT for 5 minutes, three times before the addition of secondary antibodies (Invitrogen, 1:500). Incubation was done for at least 2 hours at RT in the dark in a humidified chamber. Samples were washed three times with DPBS to remove excess secondary antibodies, each time for 5 minutes. DAPI counterstain was added with either the VECTASHIELD mounting mediumor during the secondary antibody incubation. Rectangular glass coverslips with a thickness of #1.5 were used to cover the slices.

### Quantification of SYN1/PSD95 in Cortical Organoids

Synaptic proteins SYN1 and PSD95 were quantified along the MAP2-positive area, which indicates the area occupied by the organoid slice. Images were acquired using a Zeiss LSM 900 Confocal Microscope with a 63X oil immersion objective. Laser settings and field of view size were maintained consistently throughout the imaging process. For analysis, .ics images were analyzed using an automated script from Zen Blue (Zeiss). Punctae intensity, number, and percentage of overlapping signals were measured for all channels. At least three slices per organoid were selected, with at least three images per slice used for quantifications. All values were subjected to mean.

### HiPSC-Derived Cortical Organoid Bulk RNA-Sequencing Data

Our study evaluated cortical organoid samples up to 150 days old, corresponding to human pre-natal development periods (approximately up to 19 post-conception weeks: PCW) (Gordon et al., 2021). To gain insights into the mechanisms resulting from the mutation, we performed bulk RNA sequencing (RNA-Seq) analysis on whole cortical organoids, comprising six control and six mutant samples. Cortical organoids were removed from their plate, centrifuged at 300G for 5 minutes, washed once with PBS, and resuspended in ML lysis buffer (Macherey-Nagel, Catalog No. 740971.50). One organoid was collected per tube and stored at −80 °C before RNA Extraction. To this end, we extracted total RNA from the cortical organoids using the NucleoSpin miRNA Kit for the Isolation of Small and large RNA (Macherey-Nagel, Catalog No. 740971.50) according to the manufacturer’s instructions. For cDNA amplification, we performed PCR using the Maxima First Strand cDNA Synthesis Kit according to the following protocol: HOTLID-105.20, 20 μL loading volume, 25°C for 12 min, 50°C for 15 min, 85°C for 5 min, and 4°C indefinitely (C1000 Thermal Cycler, BIO-RAD). Quantitative real-time PCR (qPCR) was performed using the Toyobo THUNDERBIRD® SYBR® qPCR Master Mix, with the following protocol: 95°C for 1:30 mins, followed by two 50x cycles of 95°C for 10 seconds, 55°C for 20 seconds, and 72°C for 30 seconds, with one 65°C cycle for 0.05s and a final exposure of 95°C for 0.5 seconds (CFX96 Real-Time PCR Detection System, BIO-RAD), and sent to Eurofins (https://www.eurofins.com) for sequencing to confirm the presence of the Nav1.2-L1342P mutation in the organoids. Subsequently, the samples were sent to the Purdue University Genomics Core at the Bindley Bioscience Center for quality control, library preparation, and sequencing. RNA quality was assessed via Qubit fluorimetry.

Raw datasets from the Element Biosciences AVITI sequencer Reads were trimmed using fastp (v0.23.2), an ultra-fast FASTQ preprocessor (Chen et al., 2018), to remove adapter sequences and low-quality bases with a score less than Q30, retaining reads with lengths greater than 50bp post-trimming for further analysis.

Trimmed Quality-controlled passed reads were aligned to the primary assembly reference of the Homo sapiens genome (GRCh38) from Ensembl release 104 using STAR Aligner version (v2.7.10b) (Dobin et al., 2013)in two-pass mode. Read alignments were assigned to featured genes using featureCounts (v2.0.1) (Liao et al., 2014)in paired-end mode and quantified on the reverse strand.

Genes exhibiting at least five reads in more than two samples were selected for Upper-Quartile normalization. For differential expression analysis between control/wild-type (WT) and L1342P mutant sample sets, WT samples were designated as the control group, while L1342P samples were treated as the experimental group. To address unwanted variation between conditions, we applied GLM regression using RUVSeq (version 1.28.0, RUVr approach) (Risso et al., 2014) to generate residuals based on the covariates of interest. This process involved identifying five factors of unwanted variation (i.e., k = 5) and incorporating them into the design matrix in edgeR (version 3.36.0)(Robinson et al., 2010), using R (version 4.1.3)(Ihaka & Gentleman, 1996). Subsequently, edgeR was employed to fit a quasi-likelihood negative binomial generalized log-linear model and conduct gene-wise statistical tests. Genes with a false discovery rate (FDR) of less than 0.05 were classified as differentially expressed.

For pathway analyses, the clusterProfiler package (version 4.10.0, R version 4.3.2) (Ihaka & Gentleman, 1996; Yu et al., 2012)and the org.Hs.eg.db package (version 3.18.0) were used to perform over-representation analyses, utilizing the filtered genes from above as the background, with a false discovery rate (FDR) control at 5%. Enrichment analyses for Gene Ontology (GO) terms and KEGG/Reactome pathways were conducted separately for differentially expressed genes. The enrichplot (version 1.23.1.992) package was used to visualize the enrichment results. Volcano plot representation was generated using ggVolcanoPlot (https://ggvolcanor.erc.monash.edu)(Mullan et al., 2021).

### Statistical Analysis

GraphPad Prism (version 9.5.1) and OriginPro 2021b (version 9.8.5.201) were used for statistical analysis. Analyses were the *Unpaired Student’s t-test, Mann-Whitney U tests,* or *One-Way ANOVA*. Results were presented as mean ± standard error of the mean (s.e.m). Values are shown in figures as *p < 0.05, **p < 0.01, ***p < 0.001, ****p < 0.0001, and n.s. (not significant) as p ≥ 0.05. The detailed statistical methods were outlined in the figure legends.

## References

Asadollahi, R., Delvendahl, I., Muff, R., Tan, G., Rodríguez, D. G., Turan, S.,…Boonsawat, P. (2023). Pathogenic SCN2A variants cause early-stage dysfunction in patient-derived neurons. Human Molecular Genetics, 32(13), 2192–2204.

Avansini, S. H., Puppo, F., Adams, J. W., Vieira, A. S., Coan, A. C., Rogerio, F.,…Montenegro, M. A. (2022). Junctional instability in neuroepithelium and network hyperexcitability in a focal cortical dysplasia human model. Brain, 145(6), 1962–1977.

Chen, G. T., Nair, G., Osorio, A. J., Holley, S. M., Ghassemzadeh, K., Gonzalez, J. G.,…Geschwind, D. (2024). Enhancer-targeted CRISPR-Activation Rescues Haploinsufficient Autism Susceptibility Genes. bioRxiv, 2024.2003. 2013.584921.

Chen, S., Zhou, Y., Chen, Y., & Gu, J. (2018). fastp: an ultra-fast all-in-one FASTQ preprocessor. Bioinformatics, 34(17), i884–i890.

Coley, A. A., & Gao, W.-J. (2019). PSD-95 deficiency disrupts PFC-associated function and behavior during neurodevelopment. Scientific Reports, 9(1), 9486.

Crawford, K., Xian, J., Helbig, K. L., Galer, P. D., Parthasarathy, S., Lewis-Smith, D.,…O’Brien, M. (2021). Computational analysis of 10,860 phenotypic annotations in individuals with SCN2A-related disorders. Genetics in Medicine, 23(7), 1263–1272.

Crowe, S. L., Tsukerman, S., Gale, K., Jorgensen, T. J., & Kondratyev, A. D. (2011). Phosphorylation of histone H2A. X as an early marker of neuronal endangerment following seizures in the adult rat brain. Journal of Neuroscience, 31(21), 7648–7656.

Dobin, A., Davis, C. A., Schlesinger, F., Drenkow, J., Zaleski, C., Jha, S.,…Gingeras, T. R. (2013). STAR: ultrafast universal RNA-seq aligner. Bioinformatics, 29(1), 15–21.

Gelot, A.-B., Courtin, T., Sileo, C., Keren, B., Soreze-Smagghue, Y., Whalen, S., & Represa, A. (2022). Polymicrogyria with Dysmorphic Neurons in a Patient with SCN2A Mutation. Journal of Neuropathology & Experimental Neurology, 81(9), 758–761.

Gordon, A., Yoon, S.-J., Tran, S. S., Makinson, C. D., Park, J. Y., Andersen, J.,…Huguenard, J. R. (2021). Long-term maturation of human cortical organoids matches key early postnatal transitions. Nature neuroscience, 24(3), 331–342.

Gould, E., & Kim, J. H. (2021). SCN2A contributes to oligodendroglia excitability and development in the mammalian brain. Cell reports, 36(10).

Hackenberg, A., Baumer, A., Sticht, H., Schmitt, B., Kroell-Seger, J., Wille, D.,…Plecko, B. (2014). Infantile epileptic encephalopathy, transient choreoathetotic movements, and hypersomnia due to a De Novo missense mutation in the SCN2A gene. Neuropediatrics, 261–264.

Hiranuma, M., Okuda, Y., Fujii, Y., Richard, J.-P., & Watanabe, T. (2024). Characterization of human iPSC-derived sensory neurons and their functional assessment using multi electrode array. Scientific Reports, 14(1), 6011.

Howell, K. B., McMahon, J. M., Carvill, G. L., Tambunan, D., Mackay, M. T., Rodriguez-Casero, V.,…Calvert, S. (2015). SCN2A encephalopathy: a major cause of epilepsy of infancy with migrating focal seizures. Neurology, 85(11), 958–966.

Ihaka, R., & Gentleman, R. (1996). R: A Language for Data Analysis and Graphics. Journal of Computational and Graphical Statistics 5: 299. doi: 10.2307/1390807.

Kang, Y., Zhou, Y., Li, Y., Han, Y., Xu, J., Niu, W.,…Huang, W. (2021). A human forebrain organoid model of fragile X syndrome exhibits altered neurogenesis and highlights new treatment strategies. Nature neuroscience, 24(10), 1377–1391.

Kearney, J. A., Yang, Y., Beyer, B., Bergren, S. K., Claes, L., DeJonghe, P., & Frankel, W. N. (2006). Severe epilepsy resulting from genetic interaction between Scn2a and Kcnq2. Human molecular genetics, 15(6), 1043–1048.

Kim, J.-E., Lee, D.-S., Wang, S. H., Kim, T.-H., & Kang, T.-C. (2024). GPx1-ERK1/2-CREB pathway regulates the distinct vulnerability of hippocampal neurons to oxidative stress via modulating mitochondrial dynamics following status epilepticus. Neuropharmacology, 260, 110135.

Kruth, K. A., Grisolano, T. M., Ahern, C. A., & Williams, A. J. (2020). SCN2A channelopathies in the autism spectrum of neuropsychiatric disorders: a role for pluripotent stem cells? Molecular autism, 11, 1–11.

Lacombe, M.-L., Tokarska-Schlattner, M., Boissan, M., & Schlattner, U. (2018). The mitochondrial nucleoside diphosphate kinase (NDPK-D/NME4), a moonlighting protein for cell homeostasis. Laboratory Investigation, 98(5), 582–588.

Liang, L., Fazel Darbandi, S., Pochareddy, S., Gulden, F. O., Gilson, M. C., Sheppard, B. K.,…Rubenstein, J. L. (2021). Developmental dynamics of voltage-gated sodium channel isoform expression in the human and mouse brain. Genome medicine, 13, 1–14.

Liao, Y., Smyth, G. K., & Shi, W. (2014). featureCounts: an efficient general purpose program for assigning sequence reads to genomic features. Bioinformatics, 30(7), 923–930.

Mao, M., Mattei, C., Rollo, B., Byars, S., Cuddy, C., Berecki, G.,…Apted, D. (2024). Distinctive in vitro phenotypes in iPSC-derived neurons from patients with gain-and loss-of-function SCN2A developmental and epileptic encephalopathy. Journal of Neuroscience, 44(8).

Matalon, D., Goldberg, E., Medne, L., & Marsh, E. D. (2014). Confirming an expanded spectrum of SCN2A mutations: a case series. Epileptic Disorders, 16(1), 13–18.

Miao, P., Tang, S., Ye, J., Wang, J., Lou, Y., Zhang, B.,…Feng, J. (2020). Electrophysiological features: the next precise step for SCN2A developmental epileptic encephalopathy. Molecular Genetics & Genomic Medicine, 8(7), e1250.

Miura, Y., Li, M.-Y., Revah, O., Yoon, S.-J., Narazaki, G., & Pașca, S. P. (2022). Engineering brain assembloids to interrogate human neural circuits. Nature protocols, 1–21.

Mullan, K. A., Bramberger, L. M., Munday, P. R., Goncalves, G., Revote, J., Mifsud, N. A.,…Purcell, A. W. (2021). ggVolcanoR: A Shiny app for customizable visualization of differential expression datasets. Computational and Structural Biotechnology Journal, 19, 5735–5740.

Parenti, I., Leitão, E., Kuechler, A., Villard, L., Goizet, C., Courdier, C.,…Bruel, A.-L. (2022). The different clinical facets of SYN1-related neurodevelopmental disorders. Frontiers in cell and developmental biology, 10, 1019715.

Pérez-Palma, E., May, P., Iqbal, S., Niestroj, L.-M., Du, J., Heyne, H. O.,…Palotie, A. (2020). Identification of pathogenic variant enriched regions across genes and gene families. Genome Research, 30(1), 62–71.

Que, Z., Olivero-Acosta, M. I., Robinson, M., Chen, I., Zhang, J., Wettschurack, K.,…Shankar, V. (2025). Human IPSC-Derived Microglia Sense and Dampen Hyperexcitability of Cortical Neurons Carrying the Epilepsy-Associated SCN2A-L1342P Mutation. Journal of Neuroscience, 45(3).

Que, Z., Olivero-Acosta, M. I., Zhang, J., Eaton, M., Tukker, A. M., Chen, X.,…Wettschurack, K. (2021). Hyperexcitability and pharmacological responsiveness of cortical neurons derived from human iPSCs carrying epilepsy-associated sodium channel Nav1. 2-L1342P genetic variant. Journal of Neuroscience, 41(49), 10194–10208.

Revah, O., Gore, F., Kelley, K. W., Andersen, J., Sakai, N., Chen, X.,…Saw, N. L. (2022). Maturation and circuit integration of transplanted human cortical organoids. Nature, 610(7931), 319–326.

Reynolds, C., King, M. D., & Gorman, K. M. (2020). The phenotypic spectrum of SCN2A-related epilepsy. European Journal of Paediatric Neurology, 24, 117–122.

Risso, D., Ngai, J., Speed, T. P., & Dudoit, S. (2014). Normalization of RNA-seq data using factor analysis of control genes or samples. Nature biotechnology, 32(9), 896–902.

Robinson, M. D., McCarthy, D. J., & Smyth, G. K. (2010). edgeR: a Bioconductor package for differential expression analysis of digital gene expression data. bioinformatics, 26(1), 139–140.

Sanders, S. J., Campbell, A. J., Cottrell, J. R., Moller, R. S., Wagner, F. F., Auldridge, A. L.,…Empfield, J. R. (2018). Progress in understanding and treating SCN2A -mediated disorders. Trends in neurosciences, 41(7), 442–456.

Sloan, S. A., Andersen, J., Pașca, A. M., Birey, F., & Pașca, S. P. (2018). Generation and assembly of human brain region-specific three-dimensional cultures. Nat Protoc, 13(9), 2062–2085. 10.1038/s41596-018-0032-7

Sloan, S. A., Darmanis, S., Huber, N., Khan, T. A., Birey, F., Caneda, C.,…Paşca, S. P. (2017). Human astrocyte maturation captured in 3D cerebral cortical spheroids derived from pluripotent stem cells. Neuron, 95(4), 779–790. e776.

Smith, R. S., & Walsh, C. A. (2020). Ion channel functions in early brain development. Trends in neurosciences, 43(2), 103–114.

Spratt, P. W., Alexander, R. P., Ben-Shalom, R., Sahagun, A., Kyoung, H., Keeshen, C. M.,…Bender, K. J. (2021). Paradoxical hyperexcitability from NaV1. 2 sodium channel loss in neocortical pyramidal cells. Cell Reports, 36(5).

Touma, M., Joshi, M., Connolly, M. C., Ellen Grant, P., Hansen, A. R., Khwaja, O.,…Agrawal, P. B. (2013). Whole genome sequencing identifies SCN2A mutation in monozygotic twins with O htahara syndrome and unique neuropathologic findings. Epilepsia, 54(5), e81–e85.

Wagner, M., Berecki, G., Fazeli, W., Nussbaum, C., Flemmer, A. W., Frizzo, S.,…Jacotin, H. (2025). Antisense oligonucleotide treatment in a preterm infant with early-onset SCN2A developmental and epileptic encephalopathy. Nature Medicine, 1–5.

Wolff, M., Johannesen, K. M., Hedrich, U. B., Masnada, S., Rubboli, G., Gardella, E.,…Villard, L. (2017). Genetic and phenotypic heterogeneity suggest therapeutic implications in SCN2A-related disorders. Brain, 140(5), 1316–1336.

Wu, W., Yao, H., Negraes, P. D., Wang, J., Trujillo, C. A., de Souza, J. S.,…Haddad, G. G. (2022). Neuronal hyperexcitability and ion channel dysfunction in CDKL5-deficiency patient iPSC-derived cortical organoids. Neurobiology of Disease, 174, 105882.

Yang, Y., Cai, Y., Wang, S., Wu, X., Shao, Z., Wang, X., & Ding, J. (2025). Human Cortical Organoids with a Novel SCN2A Variant Exhibit Hyperexcitability and Differential Responses to Anti-Seizure Compounds: Y. Yang et al.: Human Cortical Organoids with a Novel SCN2A. Neuroscience Bulletin, 1–15.

Yoon, S.-J., Elahi, L. S., Pașca, A. M., Marton, R. M., Gordon, A., Revah, O.,…Fan, H. C. (2019). Reliability of human cortical organoid generation. Nature methods, 16(1), 75–78.

Yu, G., Wang, L.-G., Han, Y., & He, Q.-Y. (2012). clusterProfiler: an R package for comparing biological themes among gene clusters. Omics: a journal of integrative biology, 16(5), 284–287.

Zhang, J., Chen, X., Eaton, M., Wu, J., Ma, Z., Lai, S.,…Lee, J. H. (2021). Severe deficiency of the voltage-gated sodium channel NaV1. 2 elevates neuronal excitability in adult mice. Cell reports, 36(5).

